# Microscopic Neuromodulation with Locally Enhanced Electric Field

**DOI:** 10.64898/2026.11.16.767056

**Authors:** Su Genelioglu, Kayeon Kim, Xu Tian, Xiyuan Liu, Changsi Cai, Anpan Han, Bingdong Chang

## Abstract

Implantable neural stimulation devices play an important role in fundamental neuroscience research, clinical applications, and emerging technologies such as neuroprosthetics and brain–machine interfaces, with increasing demand for spatially precise neuromodulation at cellular and circuit levels. Microelectrode-based electrical stimulation relies on charge injection into neural tissue, where irreversible electrochemical reactions at the electrode–electrolyte interface may lead to electrode degradation and tissue damage. Although microscale magnetic stimulation provides an alternative, it typically requires sophisticated microcoil architectures and complex fabrication processes. In this work, we introduce a novel neural stimulation mechanism based on localized electric field enhancement. The proposed device employs an electrically insulated sharp metallic tip to concentrate the electric field and generate localized displacement currents capable of activating nearby neurons. Finite element simulations were performed to investigate the underlying stimulation mechanism and establish design guidelines for device fabrication. MEMS-based neural probes were subsequently fabricated and experimentally evaluated through *in vivo* animal studies, with neuronal activation observed at distances of less than 200 µm from the probe tip using two-photon imaging. By avoiding Faradaic charge transfer at the tissue interface while offering high spatial localization, orientation selectivity, and a compact device architecture, the proposed approach provides a promising alternative to existing stimulation technologies for chronic and high-density neural interfaces.

## 1. Introduction

Neural interfaces (NIs) enable communication between the nervous system and bioelectronic devices by recording or stimulating neural activity. NIs have become a powerful tool for investigating complex neural systems, developing novel therapeutic interventions, and restoring neural function[1]. Compared with non-invasive modalities like electroencephalography (EEG)-based NIs, and transcranial magnetic stimulation [2,3], implantable NIs, including deep brain stimulation, intracortical microstimulation and microelectrode arrays [4,5], can modulate neuronal activity at cellular and circuit levels with higher spatial resolution, providing deeper insights into neural connectivity and function. Beyond fundamental research, implantable NIs have already been successfully employed in clinical settings to treat and mitigate neurological disorders like Parkinson’s disease[6], dystonia[7], essential tremor[8], epilepsy[9], and depression[9,10].

Effective neuronal modulation has been realized with different electromagnetic mechanisms. Microelectrode stimulation is efficient and typically operates at the population level. However, Faradaic current at the device-tissue interfaces can cause biotoxicity and can create challenges for chronic applications[11,12] In addition, unwanted crosstalk between closely located microprobes can interfere with independent operation[13]. Capacitive electrodes induce displacement currents without direct metal-tissue contact. However, achieving sufficient charge delivery typically requires a large electrode surface area, which limits channel density[14]. Micromagnetic stimulation is an emerging technology [15,16] and has attracted tremendous research interest due to its great potential for effective neural suppression[17]. However, the fabrication of coil-like microdevice requires advanced MEMS engineering processes[18], rendering it challenging for high-density multichannel neural interfaces. In this work, we introduce a novel electrical-field stimulation mechanism that combines high spatial precision with simplified device architectures to address these limitations.

Field enhancement effect (FEE) is a fundamental physical mechanism with broad applications across photonics, electronics, and semiconductor manufacturing[19,20]. In an electrostatic context, injecting charges into a conductive substrate within a dielectric medium causes charge density to peak at sharp geometric features. The resulting confinement of free electrons within these small volumes generates a highly localized, intensified electric field at the substrate-dielectric interface. Field enhancement factor (FEF) is intrinsically linked to the tip geometry - a principle exploited in the development of field emission devices[21]. Similar concepts have also been applied in nanophotonics to achieve local field enhancement by confining electromagnetic waves within sub-wavelength cavities for high-intensity optical resonances[22]. Inspired by the successful application of FEE in these fields, we infer that applying an alternating current (AC) to an insulated sharp tip can induce a localized polarized field, which can generate ionic currents inside neuronal tissues and effectively modulate neuronal activity. This hypothesis has been investigated systematically:

i. Numerical models were developed to investigate the mechanism and provide rational designs for microelectromechanical systems (MEMS)-based devices.
ii. The implantable devices were manufactured and packaged for good biocompatibility and minimized tissue damage.
iii. In-situ animal tests were performed on mice with two-photon microscopy (TPM) to observe neuronal responses.

The effectiveness of FEE-based neuromodulation has been successfully demonstrated with high spatial resolution. Moreover, the fabricated device has a compact design without exposed metal traces. These advantages position FEE-based neuromodulation as a promising technological platform for future NIs.

## 2. Methods

### Numerical modeling

Finite element methods (FEMs) were performed with the AC/DC module of COMSOL Multiphysics by solving the Poisson equation ∇^2^*U* = −*J_in_*/*ε* in the time domain, here *U* is the electric potential, *J_in_* is the time-varying source current density, and *ε* is the electric permittivity in the medium.

To investigate the feasibility of FEE-based neuromodulation, an ideal geometry is built in 3D cylindrical coordinates, and electrical properties are analyzed in the radial (*r)* and axial (*z)* directions. We follow the previous studies[23] on field emission devices and estimate FEF (represented by *β*) based on a hemisphere-on-orthogonal cone (hSoC) geometry. The local electric field strength ***E_e_*** can be enhanced following ***E_e_*** = *β****E*_0_**, where ***E*_0_** is the applied macroscopic electric field. For the hSoC model, FEF is given as:

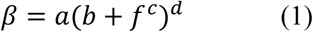

where a, b, c, and d are functions of the vertex half angle *θ*, and *f* corresponds to the apex aspect ratio *f* = *H*/*R*, here *H* and *R* represent tip length and tip radius. The displacement current generated inside brain tissues can therefore be written as:

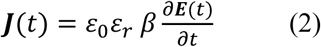

The electric displacement ***D***, which can be interpreted as charge density displaced under the polarized field, can be estimated directly as the time integral of ***J***(*t*).

For a rational design of MEMS-based devices, FEM was performed by mimicking the real device architecture, which has a 2D sharp tip geometry and is extruded in the z-direction. The electrical properties of different materials are listed in Table 1.

**Table 1.** Material properties used for numerical modeling.

|  | Aluminum | Aluminum oxide | Parylene C | Silicon | Brain tissue |
|---|---|---|---|---|---|
| Electrical conductivity (S/m) | 3.774 × 10<sup>7</sup> | 1 × 10<sup>−17</sup> | 1 × 10<sup>−14</sup> | 1 × 10<sup>-4</sup> | 0.25[24] |
| Electrical permittivity | 5000 | 5.7 | 3.2 | 11.7 | 200[25] |

### Device fabrication

MEMS-based probes were fabricated using a three-stage process starting with 100-mm-diameter, 350-µm-thick, double-side-polished (100) silicon wafers. A thermal atomic layer deposition (ALD) reactor (Picosun R200, Finland) was used to deposit a 100-nm Al₂O₃ layer using trimethylaluminium (TMA, Al(CH₃)₃; Strem Chemicals, USA) and water (H₂O) as precursors at 300 °C. A double-resist stack consisting of LOR 30C (Microresist Technology, Germany) and AZ MiR 701 (Microchemicals, Germany) was spin-coated and soft baked. Exposure was performed using a maskless aligner (MLA150, Heidelberg Instruments, Germany), followed by development in AZ 726 MIF (2.38% TMAH in water) using a puddle developer (Gamma 2M, Süss MicroTec, Germany). A 2-µm-thick Al layer was deposited by e-beam evaporation (Temescal, Ferrotec, USA), followed by lift-off in an ultrasonic bath with N-methyl-2-pyrrolidone (NMP). The LOR 30C resist layer coating was 1.2–1.3 times the thickness of the deposited Al layer to allow a clean lift-off process through undercut formation. An identical Al₂O₃ deposition step was done to fully encapsulate the Al layer. A lithography step was then performed using a 4-µm-thick AZ MiR 701 resist layer, maskless exposure, and development. Inductively coupled plasma (ICP) etching (SPTS Technologies Ltd., UK) using BCl₃/Ar plasma was employed to pattern the Al₂O₃ layer[26]. For cantilever release, a multi-step deep reactive ion etching (DRIE) process (Pegasus or ASE, SPTS Technologies Ltd., UK) was used to etch the exposed Si and define the probe geometry. The DRIE process parameters were optimized based on our previous studies[27]. For packaging, individual chips were mounted onto custom printed circuit boards (PCBs) with gold-plated bonding pads and electrically connected using Au ball wire bonding. The wire bonds were encapsulated with epoxy for mechanical protection, and the devices were conformally coated with Parylene C of varying thicknesses. Additional fabrication details are presented in the Results and Discussions section. To check the quality of insulating coatings and prevent the leakage currents, we connected the manufactured probe with a DC power supply (Keysight N5749A) and immersed the tip into phosphate buffered saline (PBS) solution (Sigma Aldrich). When sweeping the input voltage up to 100 V, no measurable current could be observed (the resolution limit of the current reading is 1 mA).

### Animal preparation and stimulation parameters

For *in vivo* validation, we used three mice (N=3), including two Thy1-GCaMP6s mice expressing a green fluorescent calcium indicator, and one mouse injected in the primary motor cortex (M1; AP +1,0 mm, ML +1.0 mm relative to bregma) with rAAV-CaMKIIa-jRGECO1a-WPRE-hGH polyA (titer: 2.48E+12v/m) two weeks prior to the experiment, enabling expression of a red fluorescent calcium indicator. Both indicators are expressed in excitatory neurons. Anesthesia was induced with 4% isoflurane and maintained at 0.9-1.5% throughout the experiment. Body temperature was continuously maintained at 37 °C. Following a craniotomy, the MEMS-based probe was inserted vertically to a depth of 200-250 µm from the cortical surface using a micromanipulator. The probe was allowed to stabilize within the tissue for at least 30 minutes before stimulation.

Stimulation was delivered through the probe connected to a function generator (Agilent 33250A). Stimulation waveforms included 1 kHz of a -90° phase-shifted (downward) half sinusoidal and a square waveform, each with a pulse duration of 1 ms. Because the stimulation paradigm is new and remains largely explorative for *in vivo* applications, we applied a broad range of stimulation intensities across experiments (0.1-10 V input voltage, 1-50 V output voltage through the signal amplifier). As the effective stimulation range varied across animals, intensities were categorized as low, medium, and high relative to the functional response range within each animal. This approach accounts for inter-animal variability likely arising from differences in probe placement, tissue-electrode coupling, local network excitability and baseline fluorescence levels. Accordingly, the minimum input voltage required to elicit a detectable neural response varied across preparations, ranging from ∼0.2 V to 2 V. All experiments were conducted using a fixed stimulation frequency of 200 Hz and a stimulation duration of 500 ms. For each condition, five trials were performed with an inter-trial interval of 10 seconds.

### TPM imaging and data analysis

Two-photon imaging was performed using a FluoView FVMPE-RS microscope (Olympus) equipped with a femtosecond laser (Mai-Tai DeepSee) and a 25× 1.05NA water-immersion objective. Imaging was conducted at an excitation wavelength of 920-980 nm, with acquisition rates ranging from 2.44 to 5.78 Hz. For data processing, image sequences were first corrected for motion artifacts using the NoRMCorre algorithm[28]. We then quantified changes in calcium fluorescence intensity within the imaged field of view by selecting regions of interest (ROI) proximal to the probe (0.1-0.2 mm), where the response peak was maximal relative to the probe location. To assess stimulation-dependent effects, response amplitudes were compared across experimental conditions, including stimulation intensity, different waveforms, probe parylene C coating thickness, and probe radius. The response amplitude (ΔF/F %) was defined as the peak response within a 3 s post-stimulation window. The baseline was calculated from the -3 to -1 s pre-simulation window. Statistical significance of differences in response magnitude between conditions was evaluated using paired or unpaired *t*-tests, as appropriate.

## 3. Results and discussions

We verified the feasibility of FEE-based neuromodulation by numerical modeling with an ideal hSoC geometry, after which a more realistic model was established to guide the design of MEMS-based implants. The device was then manufactured and packaged. Finally, *in vivo* animal tests were performed to validate FEE-based stimulation method.

### 3.1 Numerical study of FEE-based neuromodulation mechanisms

To investigate the physical mechanisms of FEE-based neuromodulation, we introduced a numerical model based on an ideal hSoC-shaped tip geometry, which is widely used for field emission devices[29]. This model enables analysis of how an applied AC input can induce current directly inside the tissue around the tip of the probe. A parameter study was further performed to understand this novel stimulation mechanism.

**Figure 1a** schematically illustrates the *in vivo* setup, where a neural implant is inserted inside a mouse brain, and TPM is employed to record the neuronal activities. Here, a “tone-arm”-shaped implant is illustrated, which has been demonstrated to have good mechanical robustness and suitable for high-resolution TPM imaging system[30]. Figure 1b presents a schematic comparison of three neuromodulation devices. To directly compare the difference in stimulation mechanisms, equivalent circuits at device-tissue interfaces are presented in Figure 1c: (i) Microelectrode stimulation relies on conductive metallic tips to introduce charges directly inside the targeted tissue. Different materials have been demonstrated, such as platinum, iridium oxide, and conductive polymers[12,31]. However, Faradaic current at the electrode-electrolyte interface is difficult to avoid, and additional measures, such as biphasic input, can be necessary to reduce the risk of electrolysis[32]. (ii) Micromagnetic simulation generates a localized magnetic field by microcoil-like device. An electric field gradient is then induced inside the brain tissue for effective neuromodulation. Since the whole device is covered by insulating layers, better biocompatibility can be achieved preferably for chronic implants. A half-turn microcoil device has been demonstrated successfully[16,17,33], while a 3D solenoid-like geometry requires more complex fabrication processes[34]. (iii) FEE-based stimulation. By fully encapsulating a sharp conductive tip with an insulating layer, Faradaic current can be avoided, and displacement current can be induced inside brain tissue by adding an AC-input on the conductive core. The interface between FEE-based probe and tissue acts like a capacitor (*C_P_*). Unlike conventional capacitive stimulation approaches, FEE-based neuromodulation doesn’t require porous or rough material to improve the charge injection capacity (CIC) or charge storage capacity (CSC)[35,36]. Instead, a sharp tip is utilized to achieve a locally enhanced electric field, that is fundamentally different to other studies on capacitor-like implant devices. Here, *I_in_* corresponds to the input current on the probe, while *R_p_* and *R_s_* are the electrical resistances of the electrode and tissue, respectively *C_dl_* and *R_f_* represent capacitance and resistance of the double layer at the interface between implants and the tissue [37].

**Figure 1.**
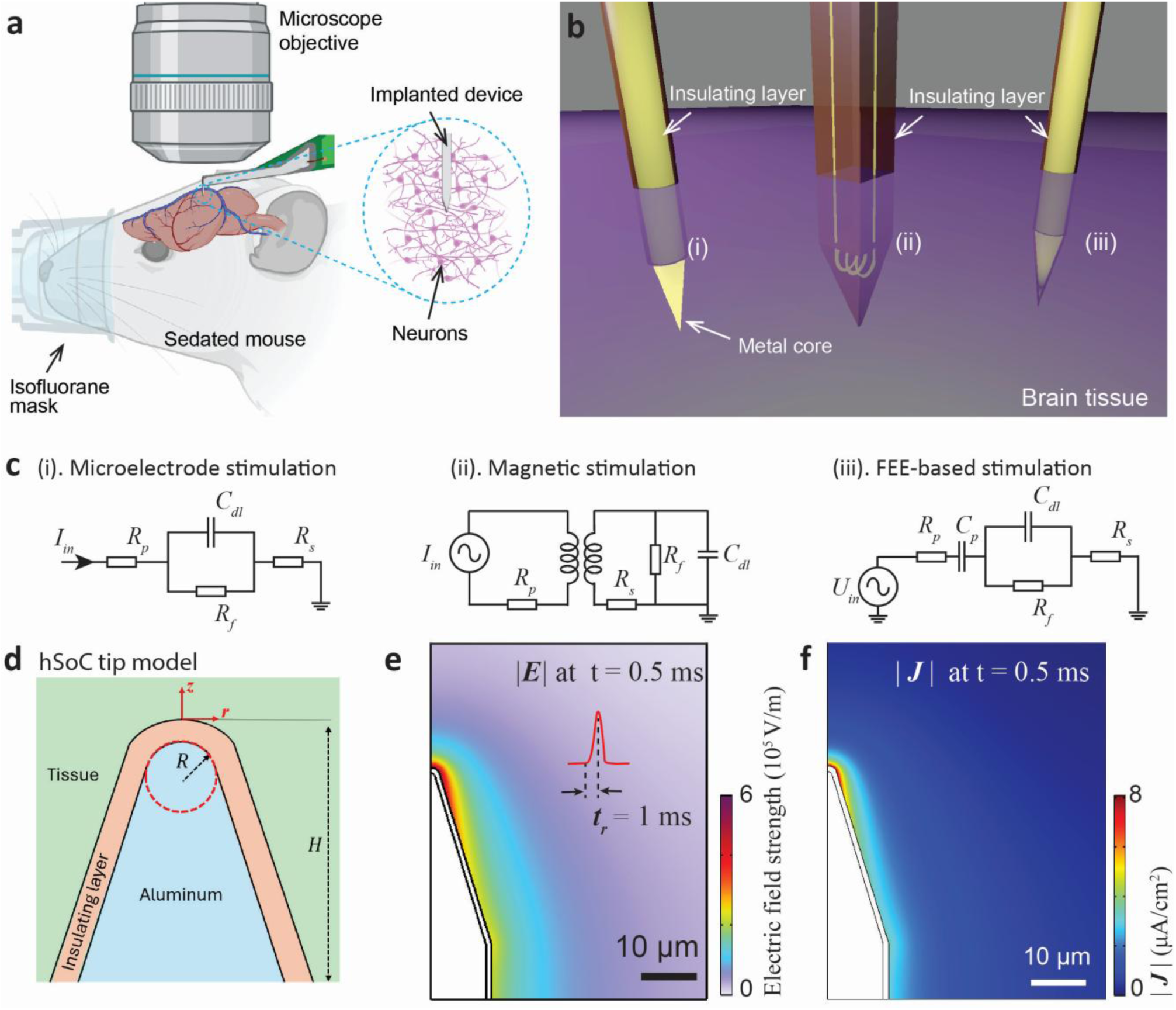
(a) Illustration of the experiment setup for an implantable device in the mouse brain, where a “tonearm”-shaped device is applied to fit the high-resolution TPM objectives. (b) Schematic drawings showing different neuromodulation implants, including (i) microelectrode, (ii) micromagnetic coil, and (iii) FEE-based probes. (c) Equivalent circuits of three neuromodulation technologies demonstrating the fundamental difference in mechanisms. (d) An ideal hSoC geometry for FEE-based neural implants. For R = 100 nm and a coating thickness of 1 µm, the distribution of electric field and displacement current are presented in (e) and (f).

Since hSoC geometry has been widely used for field emission devices, and analytic models are available to evaluate FEF, we have performed numerical modeling to verify FEE-based neuromodulations with an ideal hSoC geometry, which is illustrated in Figure 1d. We chose the tip length *H* to be 30 µm and a tip radius *R* of 1 µm, we will have apex ratio *f* = 30 and a FEF *β* ≈ 25 following equation (1). Assuming an insulating layer thickness of 2 µm and an applied voltage of 1 V with a rise time *t_r_* of 1 ms, the electric field distribution inside the tissue can then be evaluated at 0.5 ms, which is presented in Figure 1e. We can see that the normalized electric field is mostly concentrated at the tip end, with a tail extending to the lower part of the cone. The maximum electric field strength can reach around 6 × 10^5^ V/m. For purposes like field emitters, the enhanced electric field can reach the level of 10^9^ V/m[38]. The magnitude of electric displacement *D* at t 0.5 ms is presented in Figure 1f, showing a maximum current density of around 8 µA/cm^2^ close to the tip end. We also notice that the current density inside tissue decays to 50% at around 2 µm from the tip surface, showing the high locality of the stimulation process, which could be promising for high-precision neuronal stimulations. It should be mentioned that the enhanced electric field also presents inside the insulating layer, which requires sufficient dielectric strength to prevent current leakage inside the tissue. Polymer materials, like polystyrene and polyethylene, have a dielectric strength on the level of 10^7^ V/m[39], and Parylene C can have a dielectric strength of ∼ 10^8^ V/m [40], suggesting that FEE-based neuromodulation is feasible from a materials perspective.

To achieve even higher local field strength and current density inside the tissue, we keep *H* the same and reduce tip radius *R* from 3 µm to 100 nm. Based on the hSoC model, we can estimate a tip radius of 100 nm can give the highest FEF of 166, much larger than 25 (*R* = 1 µm) and 11.2 (*R* = 3 µm). The distribution of displacement current inside the tissue is compared directly in **Figure 2a**, showing a significantly increased current density with a sharper tip. The magnitude of displacement current density is plotted within 10 µm distance close to the tip end for three different tip radii (Figure 2b), and a small tip radius of 100 nm can increase the current density by at least one magnitude compared with R = 1 µm or 3 µm. For a pulse with *t_r_* = 0.5 ms and a voltage input of 10 V, the charge density can be estimated to be 0.5 µC/cm^2^, which is already above the threshold charge density for certain neuronal activation[32,41]. For a broader range of neural engineering, a higher voltage input or temporal summation can be considered. Besides, even sharper tips with *R* below 100 nm can also be technically feasible for improved charge density. Electrochemical etching can generate a tip radius of a few nanometers[42]. By applying the advanced lithography method, like electron beam lithography followed by plasma etching or lift-off, extremely sharp metallic tips can also be fabricated with high precision. We should also consider the thickness of the insulating layer, which will inevitably reduce the sharpness of the tip end. Therefore, a thinner insulation layer with a high dielectric strength will be highly favorable. This could be challenging for the coating process practically, where defects like pinholes and nonuniformity can lead to device failure. Techniques like ALD can be a promising solution, where pinhole-free insulating layers can be deposited conformally with atomic precision.

**Figure 2.**
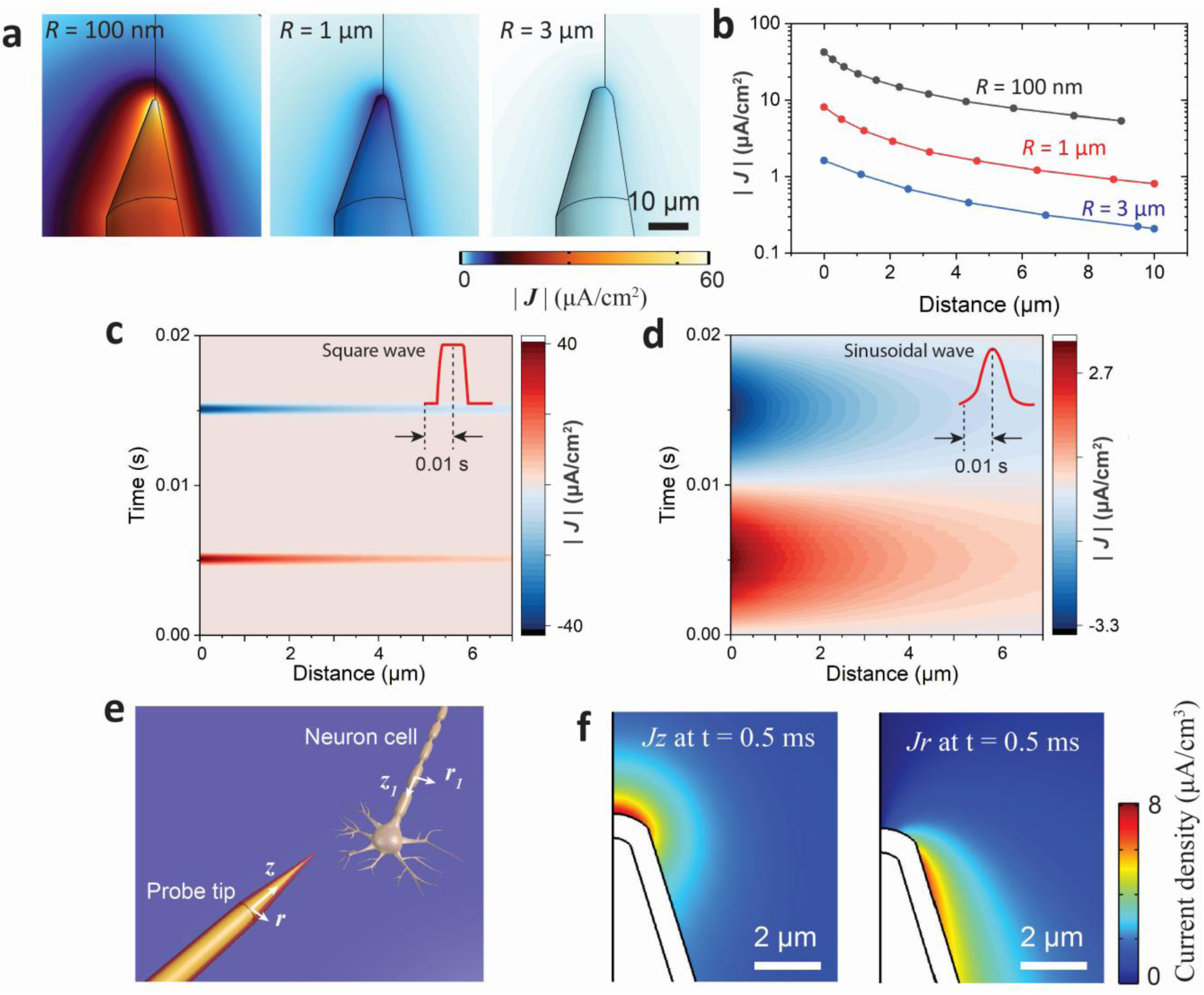
Parameter study of FEE-based neural stimulation with numerical modeling. (a) Distribution of displacement current magnitude |**J**| for different tip radii. (b) Profile of |**J**| away from the tip surface for different tip radii. (c) Time evolution of |**J**| with an input of a square waveform. (d) Time evolution of |**J**| with an input of a sinusoidal waveform. (e) Illustration of the relative position between a targeted neuron and an implanted FEE-probe. (f) Spatial distribution of J_z_ and J_r_ around the tip.

To achieve an intensified displacement current inside the tissue, another important parameter is the waveform of voltage input *U_in_*. Follow equation (2), a rapid switching of electric field is in favor. Here, we compare square and sinusoidal waveforms, which are presented in Figure 2c. Both waveforms have a full width at half maximum (FWHM) of 10 ms. For square waves, *t_r_* was set to be 1 ms, and a larger *t_r_* of 10 ms was set for sinusoidal input. The magnitude of displacement current is then plotted within a time span of 20 ms and 7 µm from the tip surface. It can be observed that a square waveform can lead to a maximum current of ∼ 40 µA/cm^2^, which is one magnitude higher than sinusoidal input (3.4 µA/cm^2^). Practically, the metallic core can be approximated as a resistor-capacitor (RC) circuit, and the input signal will be attenuated by the RC filter, resulting in a slower switching of the input signals. Therefore, high electric conductivity will be favorable for the metallic core to achieve the most efficient stimulation results. For microelectrode simulation, iridium oxide and platinum/iridium alloys are typically used, both of which have electrical conductivity on the level of 10^6^ S/m, which is one magnitude smaller than Al, it is therefore suggested that Al to be used for an optimal neuromodulation performance.

Based on numerical models, we predict another attractive feature of FEE-based neuromodulations regarding its orientation selectivity. Figure 2e illustrates the probe and a targeted neuron, the orientation of the probe and neuron can be defined by their own normalized vectors 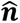 and 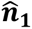 in cylindrical space (*z, r, φ*) and (*z_1_, r_1_, φ_1_*). By evaluating the distribution of displacement current density in the z- and r-components (*J_z_* and *J_r_),* we can see that *J_z_* and *J_r_* are maximized in well-separated regions around the tip. *J_z_* is mostly intensified at the very end of tip, while maximal *J_r_* is distributed away from the tip position. Since the effective extracellular activation is achieved by creating a field gradient 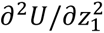 over threshold[43] for potential propagates along the axon[44], the most efficient stimulation will be achieved, when the probe tip is located close to the neuron, and 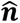 is perpendicular to 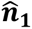. If the neuron is positioned parallel to the probe, the neuron must be located close to the sidewall of the probe to be activated. This intrinsic orientation-dependency has been demonstrated with microcoil-based magnetic stimulations[45]. For microelectrode stimulation, special designs have to be integrated to engineer the distribution of electric field for orientation selectivity[46].

Conclusively, we have demonstrated the physical mechanism for FEE-based neuromodulation with numerical methods. Unique features have been predicted regarding high locality and orientation selectivity. We also presented key parameters to be optimized for effective neuromodulation, including tip radius, thickness of insulating layer, input waveform, and electrical conductivity of metallic core.

### 3.2 Rational design of MEMS-based microprobes guided by numerical simulations

As we have numerically verified the feasibility of FEE-based neuromodulation with an ideal hSoC geometry, device architecture and design guidelines will then be established in this session. To achieve sharp metal tips with controllable tip radius, we chose to use MEMS-based fabrication technologies in semiconductor industries. A rational design was then performed with numerical modeling for effective neuromodulation.

Based on the earlier discussion, we have formulated following frameworks for numerical modeling and rational design of MEMS-based devices:

i. Al is used as the metal core for good electrical conductivity. A total thickness of 2 µm is used for Al to reduce the RC time constant.
ii. UV lithography is utilized to define 2D patterns, which can generate a reproducible resolution limit of ∼ 2 µm.
iii. A small rise time *t_r_* is evaluated for input waveforms.
iv. Al_2_O_3_ is used as a uniform coating on Al core to prevent current leakage.
v. The thickness of Parylene C coating should be relatively thin to preserve the sharp geometry at the tip.

Compared to the hSoC model, MEMS-based device is based on 2D planar Si wafers and therefore gives a tip geometry extruded in the direction perpendicular to wafer surface, instead of the ideal geometry with a rotational symmetry. To investigate FEE in a more realistic geometry, we performed numerical simulations based on a microcoil device, which has been demonstrated to have good mechanical robustness and suitable for high-resolution TPM measurements.

**Figure 3a** shows the tip design of a MEMS-based probe for FEE-based neuromodulations. An Al core is patterned on Si substrate with *W_Si_* = 70 µm, *H_Si_* = 140 µm, and *T_Si_* = 70 µm. The Al core has a thickness *T_Al_* of 2 µm. A conformal coating of Al_2_O_3_ was applied on the Al core with a thickness set to be 100 nm. Parylene C was then used to coat the whole device with a thickness of *T_P_.* Illustrative images are presented to show the material components at the end and tip of the implant device (Figure 3a (I) and (II)). The tip radius *R* of the Al trace is visualized as in Figure 3a (III). Given an Al thickness of 2 µm, we expect a minimum and reproducible *R* to be around 3 µm from a metal lift-off process, therefore we performed numerical simulations to evaluate the magnitude of displacement current, when *R* is set as 3 µm, 5 µm, and 7 µm. Figure 3b shows the distribution of |***J***| in the xz-plane inside brain tissues, when a square waveform is introduced with *t_r_* = 0.5 ms and amplitude of 50 V. We can clearly observe an enhanced displacement current density for a smaller *R*. In the *yz-*plane, FEE is also present, but the magnitude of current density is generally smaller than in the xz-plane.

**Figure 3.**
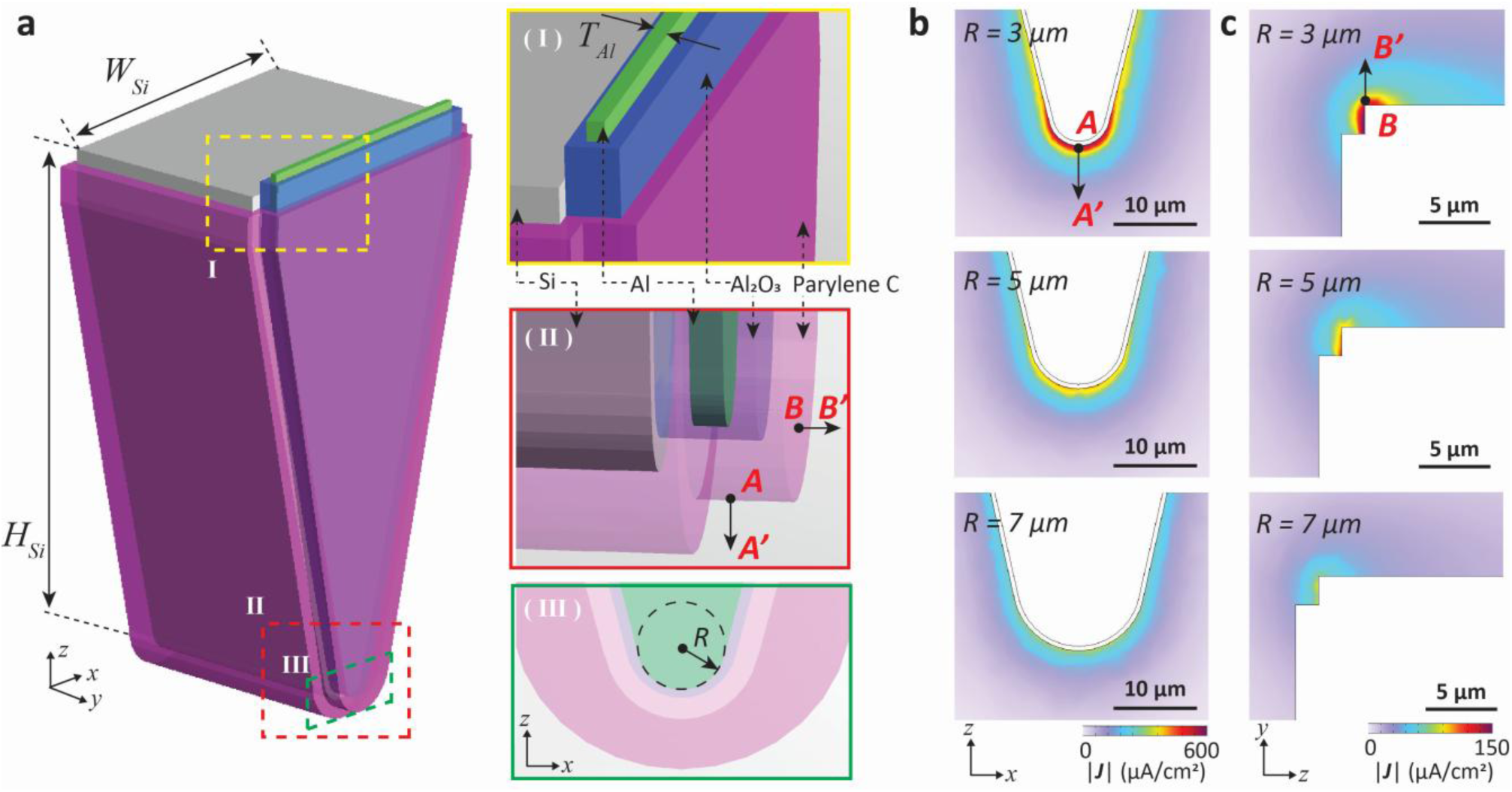
FEE with MEMS-based probes. (a) The tip of a probe showing different material components including Al core (green), Al2O3 coating (blue), Parylene C coating (purple), and Si (gray). (b) Simulated displacement current |**J**| with different tip radius in xz-plane. (c) Simulated displacement current |**J**| with different tip radius in yz-plane.

Based on the realistic model, parameter studies are performed to provide a clear specification for device manufacturing and experiment protocol. Firstly, we compared different tip radius of 3 µm, 5 µm, and 7 µm, and the displacement current density in z-direction (along AA’ as illustrated in Figure 3b), as presented in **Figure 4a**. We can see a clear FEE-related improvement of current density, and a maximum of around 200 µA/cm^2^ is present for *R* = 3 µm, which is one magnitude higher than *R* = 7 µm. Figure 4B shows current density distribution along BB’ (illustrated in Figure 3c) in the *y*-direction. No clear correlation between *R* and *J_y_* can be observed and the current density is lower than *J_z_* with a sharp geometry, which suggests a more prominent FEE in the z-direction and marks an important difference compared to the ideal geometry. We also compared different thickness of Parylene C coating of 0.5 µm, 3 µm, and 5 µm, and the current density Jz (along AA’) is presented in Figure 4c. Although a relatively thick coating is typically used in microcoil devices for safety considerations, a thin coating of 0.5 µm is desired for a high achievable current density in FEE-typed devices. Lastly, different rise time *t_r_* of 0.5 ms, 1 ms, and 2 ms are compared (Figure 4d), which again demonstrates a fast switching of voltage input to be necessary.

**Figure 4.**
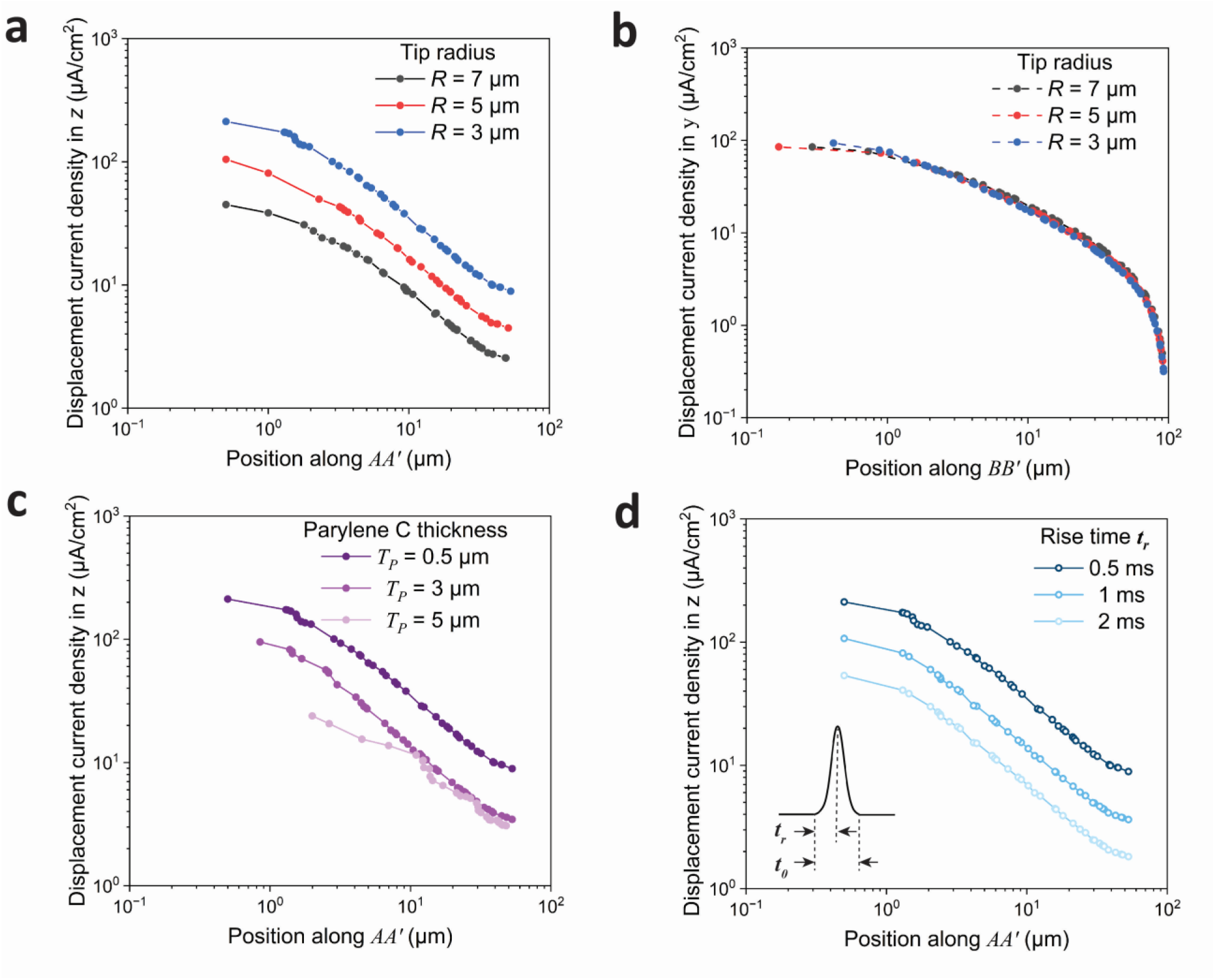
Parameter study of MEMS-based implant for neuromodulation with FEE. For different tip radius of 3 µm, 5 µm, and 7 µm, the current density distribution is presented in (a) z-direction along AA’ and (b) y-direction along BB’. (c) Current density J_z_ at the tip for different Parylene C coating thickness of 0.5 µm, 3 µm, and 5 µm. (d) Current density J_z_ at the tip when different rise time t_r_ is used for the voltage input.

To summarize, we have demonstrated the feasibility of FEE-based neuromodulation with a realistic model. Compared to the ideal geometry, the realistic model is built on a planar MEMS-based device and FEE is maximized on the in-plane surface. Our results also provide design parameters for MEMS-based implant devices to be manufactured.

### 3.3 Device fabrication

Based on the design guidelines discussed in 3.2, MEMS-based manufacturing technologies were used to develop probes for further animal tests. The fabrication process consisted of three masks and three main stages. The first stage involved patterning the metallic wire using UV lithography and thin film deposition, followed by an Al_2_O_3_ patterning stage to define the wire-bonding pads and establish the probe geometry, and the final stage involved dry Si etching for cantilever release and packaging.

The fabrication process is illustrated in **Figure 5a**, which starts with a 100-mm-diameter, 350-μm-thick, double-side-polished Si wafers. Initially, a 100-nm-layer of Al_2_O_3_ was deposited using ALD on both sides for proper insulation (i). A double-stack resist layer lithography step was then carried out for metallic wire patterning (ii). A 2-µm-thick Al layer was deposited using an e-beam evaporator, followed by a lift-off step (iii and iv). Subsequently, a second 100-nm Al_2_O_3_ layer was deposited to act as an insulation barrier as well as a hard mask during Si etching for cantilever release (vi). The Al_2_O_3_ layers were patterned using ICP etching to define the wire bonding pads and the silicon probe geometry on both sides of the wafer (vii).

**Figure 5.**
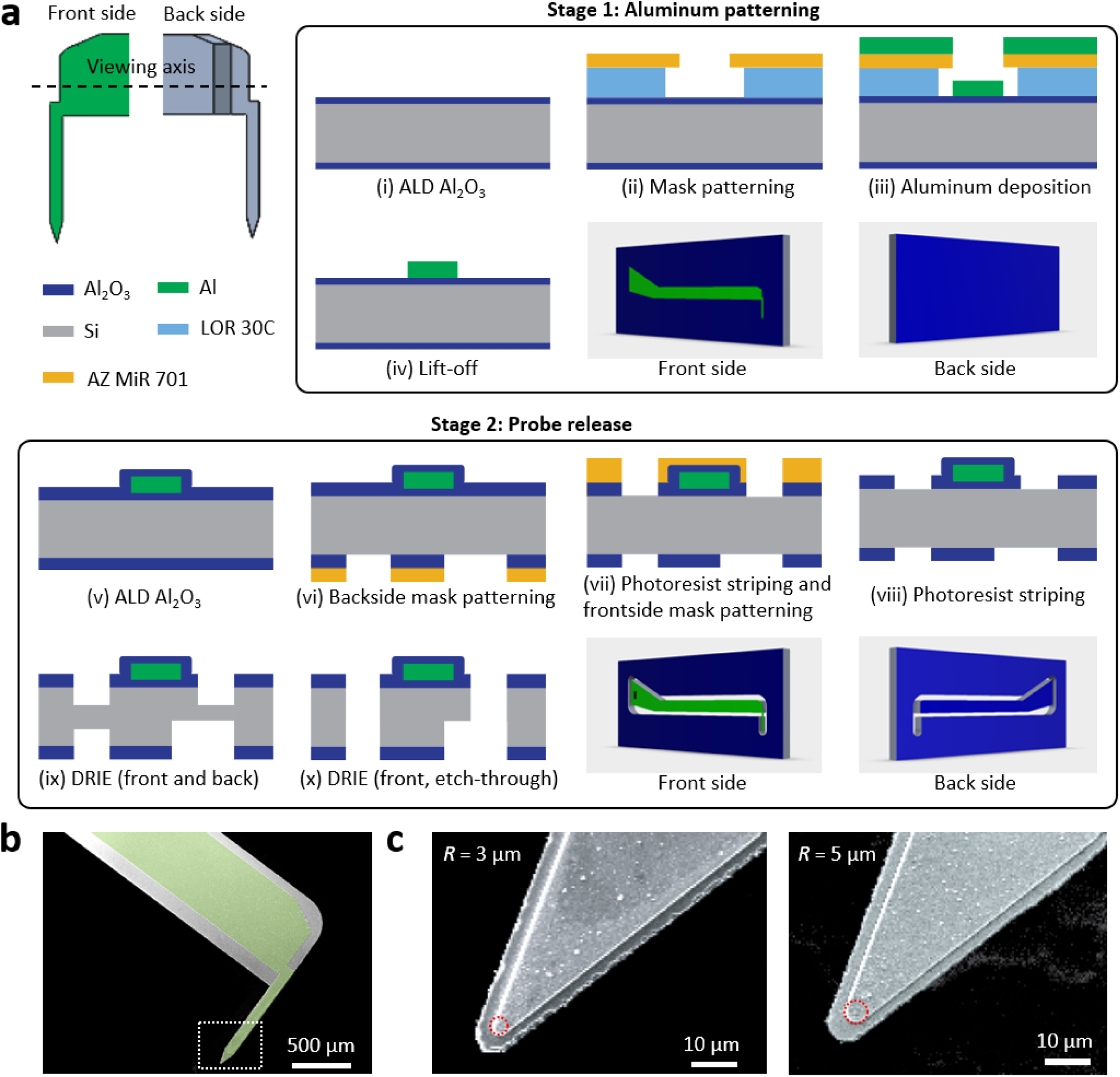
Fabrication of MEMS-based implants for neuromodulation with FEE. (a) Process flow. (b) SEM image of the cantilever and tip region, false color is used to highlight the Al trace. (c) SEM images of probe tips with R = 3 µm and 5 µm, red dash circle is used to highlight the difference in tip radius.

Finally, the photoresist residues were removed (viii) by plasma ashing and MEMS devices were released from the substrate using a multiple-step DRIE process (ix and x). During this step, a height difference was introduced between the probe tip and the remaining probe structure to minimize the tip thickness and reduce implantation-induced tissue damage. The tip of fabricated “tonearm”-like device is presented by SEM image in Figure 5b, showing both cantilever structure and the tip region to be implanted. Two different tip radius of 3 µm and 5 µm are also manufactured for direct comparison in animal tests, and the high-magnification SEM images are presented in Figure 5c.

In the final packaging stage, individual devices were then glued to a PCB and electrically connected using Au ball wire-bonding. To protect the electrical connections during animal tests, the wire bonds were covered with epoxy. The devices were then completely encapsulated with Parylene C for biocompatibility and electrical insulation. The integrity of the Parylene C coating was assessed by immersing the probes in phosphate-buffered saline (PBS) and measuring leakage current under an applied voltage using a power supply up to 50 V.

### 3.4 Animal test

To verify the physical mechanism and manufactured MEMS-based microprobes, we performed animal tests on Thy1-GCaMP6s mice and mice injected with rAAV-CaMKIIa-jRGECO1a, both expressing calcium indicators in excitatory neurons for *in vivo* validation.

**Figure 5a** shows the experiment setup for *in vivo* TPM. The input waveform is illustrated in Figure 5b, where pulse width was set as 1 ms with an inter-pulse interval of 5 ms and total number of 100 pulses in each pulse train. A schematic drawing is presented in Figure 5c to show the cranial window on the head of the mouse, and a representative two-photon image shows a clear neuronal response upon stimulation. Figure 5d shows the neuronal calcium activity of an example cell (red circle), where the response amplitude increased with stimulation intensity from low to high (see M*ethods* for details), and reached the largest response at high intensity, indicating that response is stimulation intensity-dependent since we used the same probe across these stimulation intensities applied. We observed the largest calcium increase at an approximately 100∼150 µm from the probe location, with responses becoming more apparent from medium stimulation intensity (Figure 5e). This spatial pattern may reflect local tissue effects associated with acute probe insertion or the spatial distribution of the induced field with nonlinearity.

To assess population-level activity, we analyzed the recorded field of view and quantified responses across stimulations conditions. Stimulation-evoked responses increased with input intensity for both sinusoidal and square waveforms (Figure 6f), indicating a robust intensity-dependent modulation of population activity (sinusoidal waveform in light green; low vs. medium; *p* = 0.002; low vs. high; *p* = 0.0036; medium vs. high; *p* = 0.06). Square wave stimulation similarly exhibited intensity-dependent differences across all levels (Fig. 6f, dark green; low vs. medium; *p* = 0.01; low vs. high; *p* = 3.4 × 10⁻⁴; medium vs. high; *p* = 6.8 × 10⁻⁴). Across all stimulation intensities, square waveform elicited larger responses than sinusoidal waveform, with the difference reaching statistical significance at high intensity (Fig. 6f, *p* = 0.007). These results indicate that while both waveforms drive evoked neural responses, square waveform is more effective at eliciting stronger population activity than sinusoidal waveform, consistent with our simulation results.

**Figure 6.**
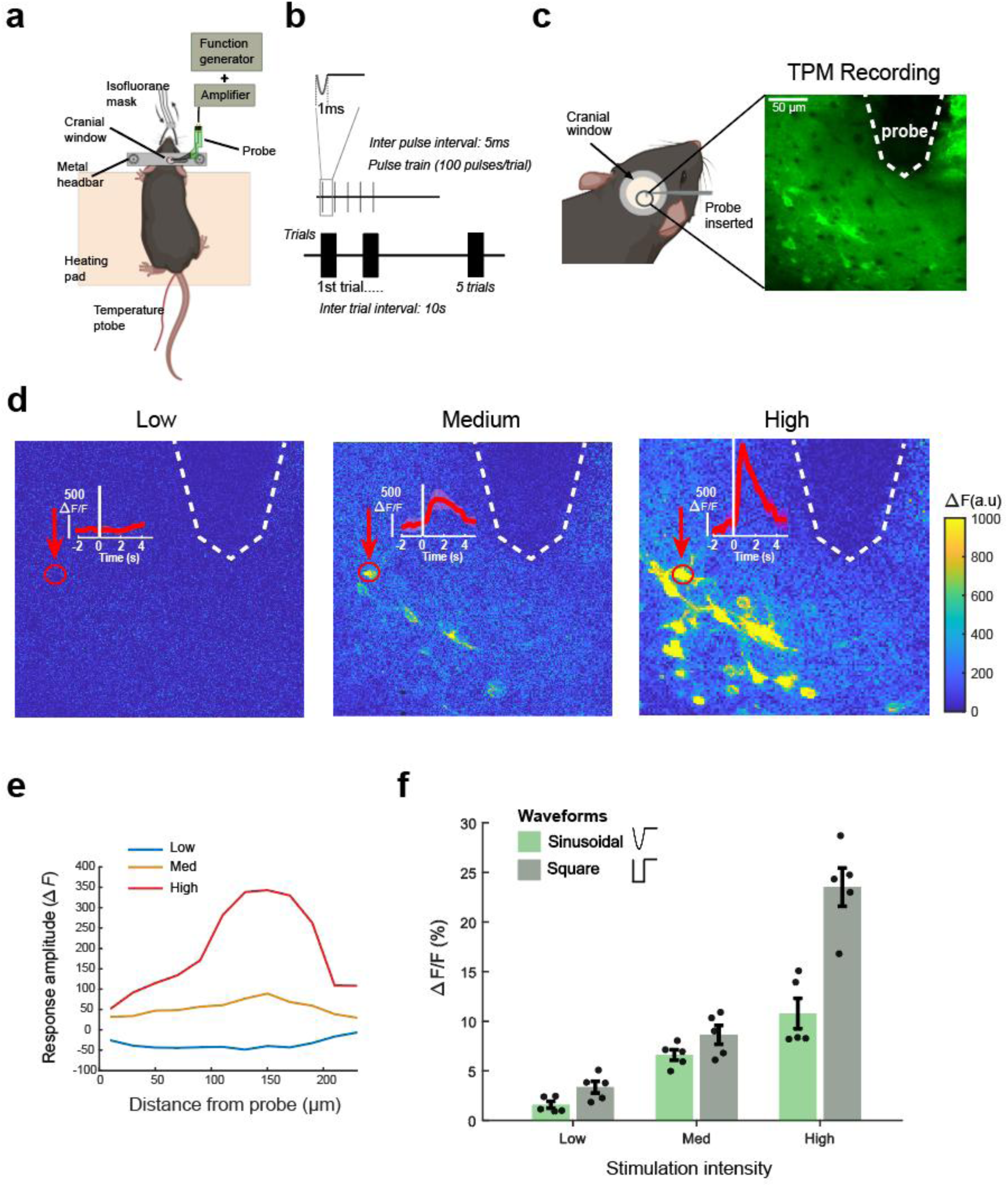
In vivo TPM experiments reveals stimulation-dependent increases in neuronal calcium responses. (a). In vivo experimental setup. The probe was mounted on a micromanipulator and connected to a function generator and power supply. (b). Stimulation parameter showing the waveform, inter-pulse interval, and inter-trial interval used in the experiments. (c). Left: Illustration of a cranial window over the mouse motor cortex, with the probe inserted 150∼200 µm below the cortical surface. Right: Representative two-photon imaging field of view showing the probe location (dashed line) and neuronal somata in the field of view. (d). Baseline-subtracted calcium response (ΔF) across the imaged field of view, with frames averaged over the 3 second following stimulation. Dashed lines indicate the probe location, and the red circle marks an example single neuron showing a stimulation-associated response. Inset: response trace aligned to stimulation timing, averaged across five repetitions using sinusoidal waveform Shaded region indicates mean ± 2 x standard error of the mean (s.e.m). (e). Response amplitude plotted as a function of distance from the probe location. (f). Bar plots showing response amplitude (ΔF/F %) as a function of stimulation intensities with mean ± 2 s.e.m. Light green represents sinusoidal stimulation, and dark green represents square wave stimulation.

To further validate the mechanism of FEE-based neuromodulation, we compared different probe design configurations. Firstly, a 0.5 µm-thick parylene C coating generate a more significant response (∼ 5 times) compared to the device with 1 µm-coating (Fig. 6a). Secondly, probe with a smaller tip radius (*R* = 3 µm) elicited larger responses than probe with 5 µm tip radius under the same stimulation intensity and coating thickness (Figure 7b). These observations agree nicely with our predictions from numerical modeling.

**Figure 7.**
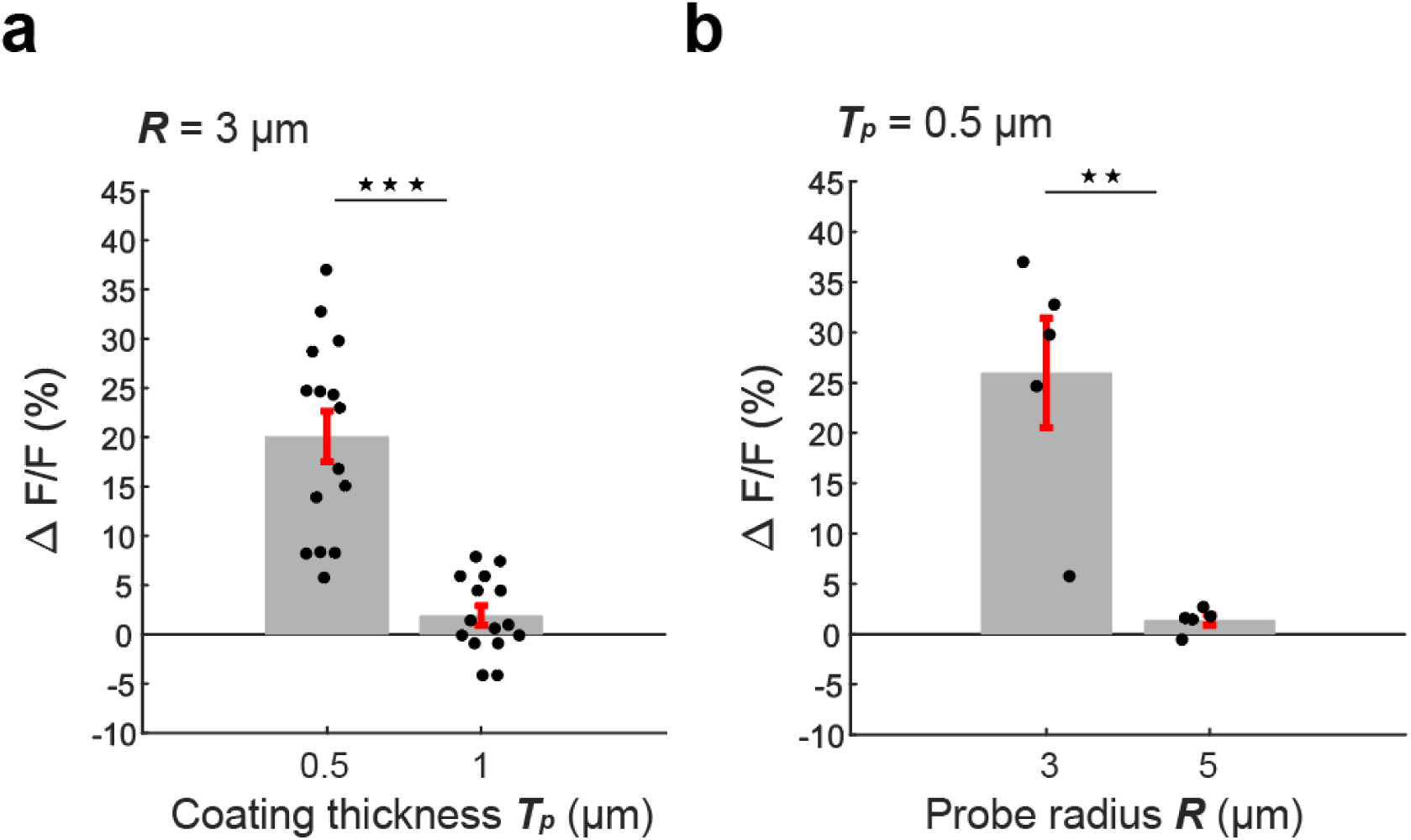
Influence of Parylene C thickness and tip radius on stimulation-associated calcium responses. (a) Response amplitude plotted as a function of parylene coating thickness, with probe tip radius and input voltage held constant. (b) Response amplitude plotted as a function of probe tip radius, with coating thickness and input voltage (at high level) held constant. Bar represent mean ± 2 s.e.m. ***p < 0.001, ** p < 0.01

Overall, these *in vivo* observations provide supporting evidence that the proposed stimulation approach can modulate neuronal calcium activity in the living mouse brain. While further experiments are needed to fully characterize the cellular mechanisms, these results warrant future investigation and optimization of this stimulation method.

## 4. Conclusions

To summarize, we propose a novel strategy for neural stimulation. Physical models have been developed, and numerical simulations have been performed to verify the theory and to provide guidelines for device fabrication. Compared with other stimulation methods like microelectrodes, capacitor electrode, and micro-coils, this strategy is based on FEE at a sharp probe tip, which can directly induce displacement current flux inside tissue. The device geometry and material components mark FEE-based stimulation to be fundamentally different compared to other NI technologies. Although FEE has been noticed and utilized to improve the performance of other types of neural stimulation devices, like micro-coils[47,48], our study marks the first attempt to use FEE directly for effective neuromodulations. Some attractive advantages of this technology are highlighted:

i. A compact device for minimal tissue damage and possibility of high-density multichannel integration.
ii. High locality with an activation area below 200 µm validated with *in vivo* experiments, which is favorable for high-resolution stimulations.
iii. Orientation selectivity as predicted by numerical modeling.

With these promising results, future work will be multi-channel integration for *in vivo* and chronic applications. Moreover, we expect *in vitro* experiments to provide insights in biophysical mechanisms, especially by modifying the frequency and polarity of input signals. We expect this work to pave the way for new possibilities in neural engineering.

## Acknowledgement

The authors would like to thank the valuable discussion with Shelley Fried, Kyeong Jae Lee, and Jae-Ik Lee from Havard Medical School.

## Author declaration

The authors declare no competing interests.

